# GRIDSS: sensitive and specific genomic rearrangement detection using positional de Bruijn graph assembly

**DOI:** 10.1101/110387

**Authors:** Daniel L Cameron, Jan Schroeder, Jocelyn Sietsma Penington, Hongdo Do, Ramyar Molania, Alexander Dobrovic, Terence P Speed, Anthony T Papenfuss

## Abstract

The identification of genomic rearrangements, particularly in cancers, with high sensitivity and specificity using massively parallel sequencing remains a major challenge. Here, we describe the Genome Rearrangement IDentification Software Suite (GRIDSS), a high-speed structural variant (SV) caller that performs efficient genome-wide break-end assembly prior to variant calling using a novel positional de Bruijn graph assembler. By combining assembly, split read and read pair evidence using a probabilistic scoring, GRIDSS achieves high sensitivity and specificity on simulated, cell line and patient tumour data, recently winning SV sub-challenge #5 of the ICGC-TCGA DREAM Somatic Mutation Calling Challenge. On human cell line data, GRIDSS halves the false discovery rate compared to other recent methods. GRIDSS identifies non-template sequence insertions, micro-homologies and large imperfect homologies, and supports multi-sample analysis. GRIDSS is freely available at https://github.com/PapenfussLab/gridss.

## INTRODUCTION

Structural variants (SVs), including large deletions and chromosomal translocations, play a significant role in the development of cancer and other diseases. Historically, early links between SVs and cancer were found by cytogenetics [e.g. 1], including the discovery of circular chromosomes [2] and the Philadelphia chromosome [3, 4]. It was only with the advent of high-throughput sequencing that the extent of genomic rearrangement in cancer, the role of catastrophic chromosome-scale events such as chromothripsis [5], the broader clinical relevance of SVs [6] and their predictive value [7] were identified. Although significant progress has been made, SVs remain less well studied than single nucleotide variation, in part due to challenges in their reliable identification from short-read sequencing data.

Many methods exist to identify SVs using high-throughput sequencing data. These use one or more of three forms of evidence: read depth, split reads, and discordantly-aligned read pairs. Changes in read depth (RD) are associated with copy number variants and imply genomic rearrangements, but RD methods [e.g. 8, 9] cannot resolve genomic fusion partners and breakpoint positions are imprecise. Rather, resolution is dependent on the overall sequencing depth and the selected window size used in the analysis. Using paired-end sequencing, clusters of discordantly-aligned read pairs (DPs)—i.e. read pairs that align to different chromosomes, with unexpected orientation, or separation—can be used to infer the presence of SVs. Since the SVs are assumed to occur in the unsequenced part of the DNA fragments, DP methods [e.g. 10, 11] do not identify exact breakpoint locations. Single nucleotide resolution of SVs is useful for predicting possible fusion gene products or the impact of a promoter translocation, identifying disrupted tumour suppressors, determining the DNA repair mechanism responsible for the SV, and investigating motifs associated with the breakpoint. It is obtained using split reads (SR), where the sequenced reads span the breakpoint. SR methods [e.g. 12, 13, 14] find SVs by identifying split alignments in which part of the read aligns to either side of a genomic rearrangement, either through direct split read mapping by read aligner, re-alignment of soft clipped (SC) bases (unaligned bases in partially mapped reads), or split alignment of the unmapped read in one-ended anchored (OEA) read pairs (read pairs with only one read mapped) [15]. Whilst most tools consider only the best alignment for each read, some methods [11, 16] consider multiple potential alignment locations to better handle repetitive sequence [17]. Multiple approaches have been developed that incorporate two (e.g. DELLY [18]) or three (e.g. LUMPY [19]) forms of evidence in the variant calling process. Long reads are now also being used [20] or incorporated as evidence (e.g. Parliament [21]).

To improve SV calls, short read assembly has also been incorporated into methods in a variety of ways. Assembly of reads obtained from clusters of SC reads (e.g. CREST [13]) or Open End Anchors (OEAs) (e.g. NovelSeq [22]) has been used to form break-end contigs, which extend out and span the breakpoint from each side. In contrast, breakpoint contigs are generated by local assembly of all reads supporting a rearrangement, generating a single contig supporting the variant. Some methods apply targeted assembly to validate the breakpoint calls (e.g. Manta [23], SVMerge [24], TIGRA [25]). Windowed breakpoint assembly has been used (e.g. SOAPindel [26], DISCOVAR [27]), but detection is limited to events smaller than the window size. Whole genome *de novo* assembly has also been used for variant calling (e.g. cortex_var [28]), but its use has been limited, in part due to the computational expense compared to alignment-based approaches.

Here, we present a novel approach to predicting genomic rearrangements from DNA sequencing data, GRIDSS (Genome Rearrangement IDentification Software Suite), which provides high-speed variant calling from a combination of assembly, split read and read pair support. The philosophy underpinning GRIDSS is to maximise sensitivity and prioritize calls into high or low confidence, thereby maintaining specificity in the high confidence call set. To achieve this, we take a three-step approach. Firstly, we filter out reads that align properly; i.e. we extract all reads that might provide any evidence for underlying genomic rearrangements. Secondly, we perform assembly of all remaining reads using a novel algorithm that utilises information from the alignment to constrain the assembly. We term this genome-wide break-end assembly, as each contig corresponds to a break-end and only after assembly is the underlying breakpoint and partner break-end identified. Unlike existing break-end assemblers that perform targeted assembly of soft clipped (SC) or one-end anchored (OEA) reads, our approach performs genome-wide assembly of all SC, SR, DP, OEA and indel-containing reads. Similar to split read identification from soft clipped reads, breakpoints are identified by realignment of break-end contigs. Finally, we apply a probabilistic model that combines break-end contigs from each side of the rearrangement with SR and DP evidence to score and call variants.

To perform the genome-wide break-end assembly, we developed a novel assembly approach specifically for the task by extending a positional de Bruijn graph data structure. Originally developed for small indel and base calling error correction of *de novo* assembly contigs [29], positional de Bruijn graphs add positional information to each node, transforming them into a directed acyclic graph and making use of valuable information generated by the aligner. With appropriate optimisation, this is computationally efficient at the genome scale, and reduces depth of coverage needed and memory requirements for accurate assembly. To make the best use of data from related samples, sequencing libraries are tracked in the de Bruijn graph using color [28], and evidence supporting rearrangements is shared between libraries during assembly and variant calling. Since the assembled contigs are longer than the read length, this improves performance in regions of poor mappability. Meaningfully scored variants and a set of useful default filters makes GRIDSS easy to use, but also a powerful tool for advanced users, who, armed with prior knowledge about expected rearrangements, can identify relevant calls with low support.

Using simulated and cell line data, we show that GRIDSS outperforms other methods. We demonstrate its effectiveness on cell line and patient tumour sequence data, including rearrangements with extreme complexity, as well as sequencing data from the AT-rich *Plasmodium falciparum* genome.

## RESULTS

### Outline of GRIDSS

The GRIDSS pipeline comprises three distinct stages: extraction, assembly, and variant calling (Figure 1).

**Figure 1.**
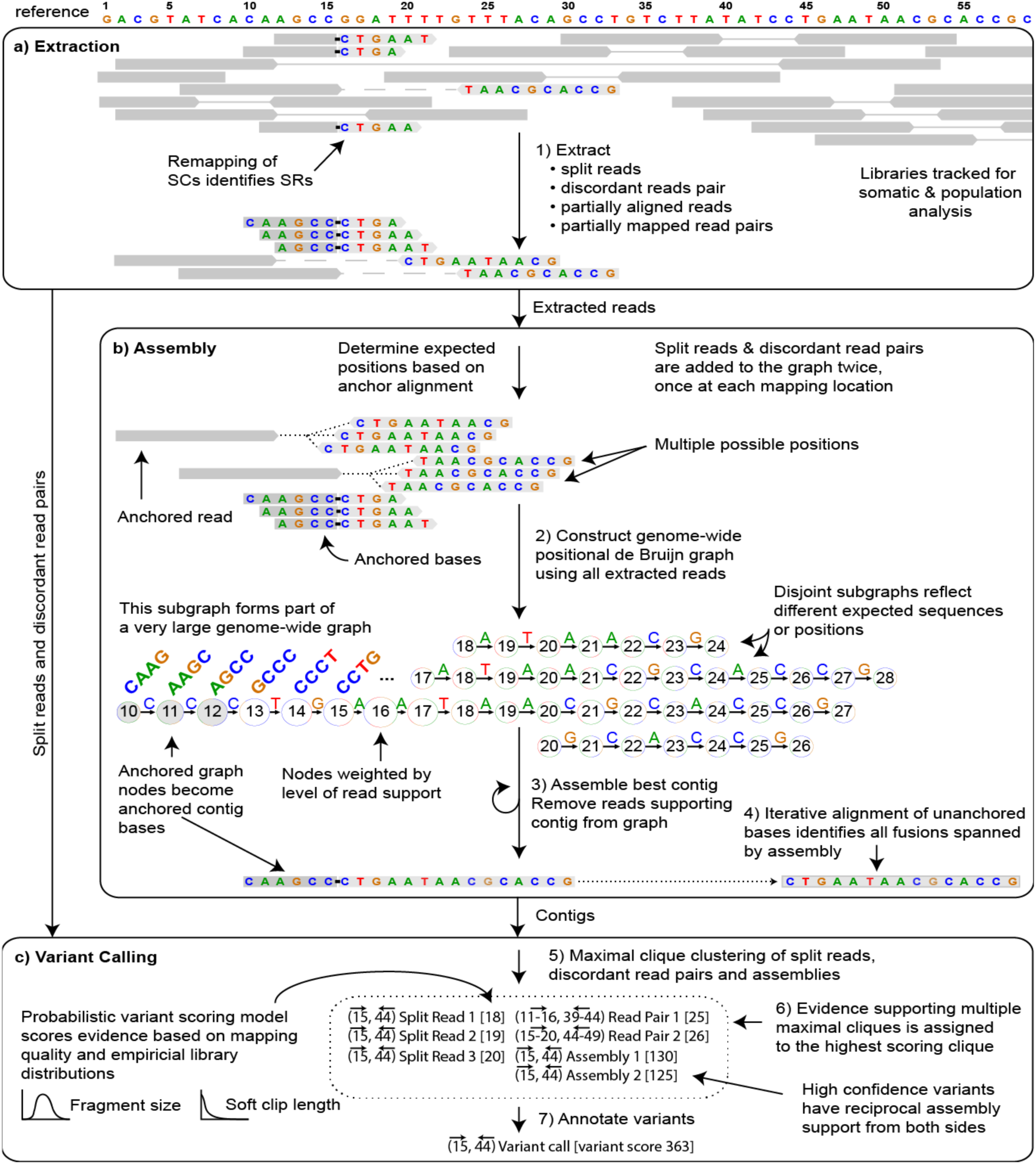
Outline of the GRIDSS pipeline. a) Soft clipped and indel-containing reads as well as discordant and one-ended anchored read pairs are extracted from input BAM files. Split reads are identified through re-alignment of soft clipped read bases. b) Extracted reads are added to a positional de Bruijn graph in all positions consistent with an anchoring alignment. Break-end contigs are identified by iterative identification of the highest weighted unanchored graph path followed by removal of supporting reads. Unanchored contig bases are aligned to the reference genome to identify all breakpoints spanned by the assembly. c) Variants are called from assembly, split read, and read pair evidence using a probabilistic model to score and prioritise variants.

In the extraction stage, GRIDSS takes as input any number of SAM/BAM alignment files and calculates a number of summary statistics for each sequencing library. SRs are identified by realignment of the unaligned 3’ or 5’ “soft clipped” read bases back to the reference genome. DPs consist of read pairs aligned with inferred fragment size shorter or longer than 99.5% of read pairs, in the wrong orientation, or to different chromosomes. All SC, DP, OEA and indel-containing reads are then extracted. At this stage no SV calling has been undertaken. Reads are scored according to likelihood of originating from the reference allele based on the read mapping quality, the empirical distributions of the read alignment and library fragment size, and read mapping rate. Extracted reads are passed to the assembly stage, with split reads and discordant read pairs also passed to the variant calling stage.

In the assembly stage, reads are decomposed into a sequence kmer of k consecutive bases. These kmers and their genomic locations are incorporated into a positional de Bruijn graph. Kmers are considered anchored if the originating read aligns to the reference for all k bases and these anchoring kmers are used to constrain the positions in which a read can be assembled. Unanchored kmers from soft clipped reads are placed as if the read were fully mapped. Split and indel-containing reads are treated as two independent soft clipped reads—one for each alignment location. OEA read kmers are placed at all positions compatible with the alignment of the mapped read and the DNA fragment size distribution, with DPs treated as two independent OEA read pairs. Each read kmer is weighted according to the constituent base quality scores and variant support score, with graph nodes weighted by the cumulative supporting weights. Error correction is performed to remove spurious paths caused by sequencing errors. Break-end contigs are iteratively identified by finding the highest weighted unanchored path and extending into anchoring kmers if present. To ensure each read supports only a single contig, reads supporting each break-end contig are removed from the graph when the contig is assembled. Unaligned contig bases are iteratively aligned to the reference genome to identify all genomic rearrangements spanned by the assembly. Data streaming and graph compression is used extensively to keep the assembly memory footprint below 2GB per thread.

Variant calling occurs in the final stage of GRIDSS. Variants are identified from the overlap in predicted breakpoint positions of assemblies, SRs and DPs. After identifying and scoring all overlapping support sets, each SR, DP and assembly is then assigned to the highest scoring variant it supports. High-scoring variants with assembly support from both break-ends are considered high confidence calls. Variants that are low-scoring or lack assembly support from one or both break-ends are considered low confidence and a filter flag is applied to these variants in the output VCF. Non-template sequence insertions as well as exact micro-homologies and large imperfect homologies are automatically identified in the variant calls.

### Performance on simulated data

To assess the performance of GRIDSS, we simulated heterozygous structural variants with a range of event types (deletion, insertion, inversion, tandem duplication, translocation) and sizes (1 base pair (bp) to 65 kilo-base pairs) on human chromosome 12 (hg19). We compared the GRIDSS results to a number of other tools (BreakDancer, Pindel, DELLY, HYDRA_MULTI, LUMPY, SOCRATES, cortex, and TIGRA; see Supplementary Material for full details).

For parameters typical of tumour genome sequencing (60x coverage of 100bp paired end reads with a mean fragment size of 300bp), GRIDSS obtained near-perfect sensitivity across the widest range of event types and sizes (Figure 2), albeit with Pindel having greater sensitivity on small (<50bp) events, and only the *de novo* assembly-based caller cortex able to detect large insertions. For both random breakpoints and breakpoints in SINE/ALU elements, only the multi-mapping aware caller HYDRA obtained sensitivity higher than GRIDSS (99.58/99.57% vs 99.36/99.22%), with DELLY, Socrates and LUMPY exceeding 95% sensitivity in both data sets (99.08/98.76%, 98.94/98.96%, 97.43/97.78%). BreakDancer, DELLY, Pindel, and TIGRA all incorrectly classified breakpoint events as inversion events. Except for the detection of insertion events using cortex, GRIDSS obtained a higher precision than other callers with comparable sensitivity (Supplementary Figure 2).

**Figure 2.**
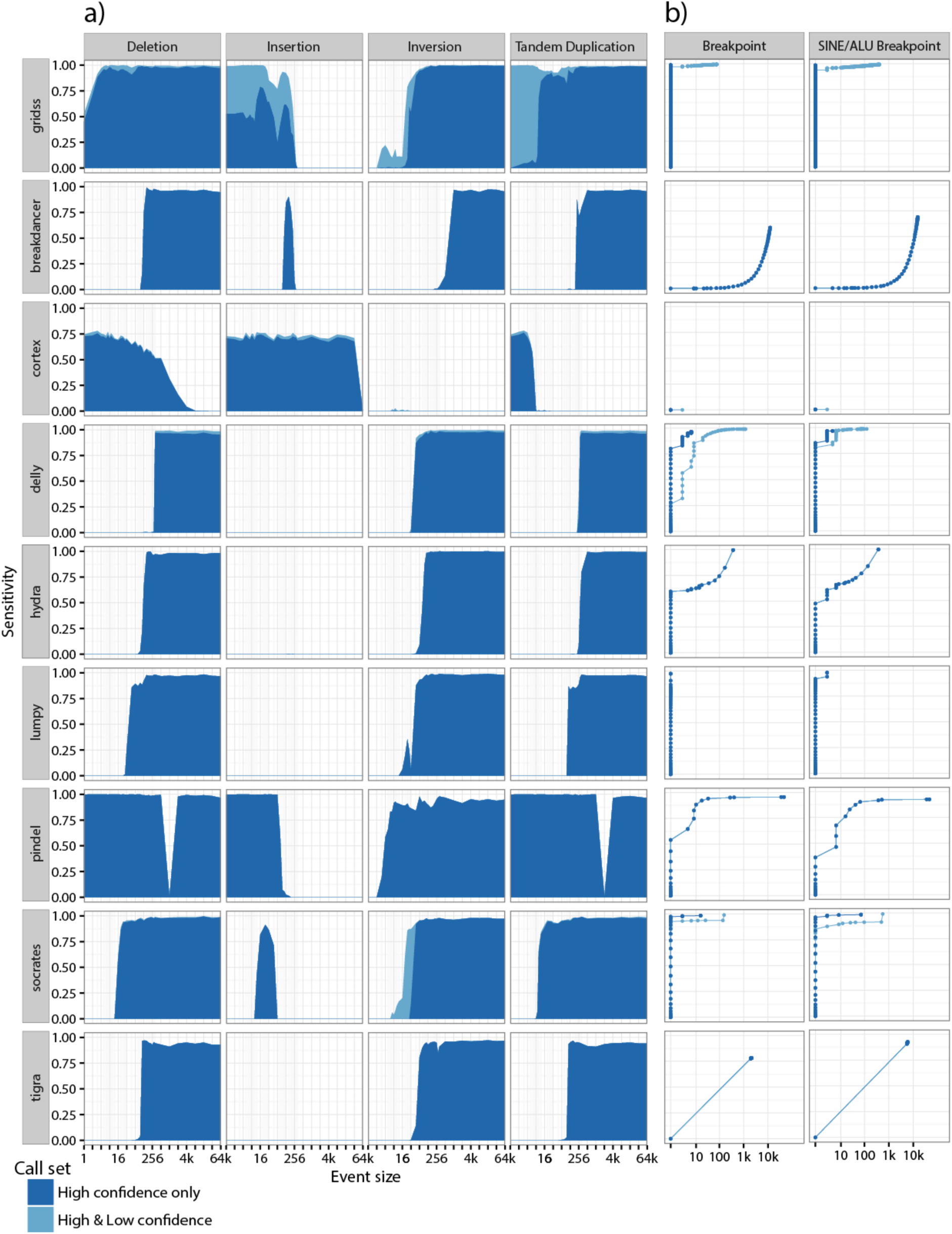
Variant caller performance on simulated heterozygous genomic rearrangements. Different classes of genomic rearrangement were randomly generated against human chr12 (hg19) and 60x coverage of 2x100bp sequencing data was simulated. (A) The sensitivity of each method (rows) for each event type (columns) is plotted against event size. (B) Receiver Operating Characteristic (ROC) curves for all breakpoints (left) and breakpoints located in SINE/ALUs (right).

To explore the impact of read length (36-250bp), DNA fragment size (100-500bp), sequencing depth (4-100x) and aligner (bwa, novoalign, bowtie2), GRIDSS was applied to a comprehensive simulation (see Supplementary Material for further details). For reads 50bp or longer, GRIDSS is able to reliably call and assemble heterozygous genomic fusions at 30x coverage regardless of aligner or fragment size, although some libraries (such as 100bp paired end reads with 500bp fragment size) require as little as 8x coverage. Overall, the F_1_-scores of GRIDSS show improved call quality for increasing read length, read depth, and fragment size (Supplementary Figure 3). Whilst the precision of calls supported by single-sided or no assembly decreased with coverage as expected, precision of calls supported by reciprocal breakpoint assembly remained near 100% regardless of sequencing depth, read length, library fragment size, or aligner (Supplementary Figure 5). This demonstrates that requiring reciprocal break-end assembly support is a simple yet surprisingly powerful false positive filter. Although frequently overlooked or uncontrolled in experimental designs, our simulation results reveal the significant impact library fragment size has on structural variant calling. Unlike single nucleotide and indel calling, which are relatively independent of library fragment size, the impact of library fragment size on structural variant calling can be the equivalent of up to a two-fold change in coverage.

### Performance on cell line data

We next tested GRIDSS on several real sequencing data sets. First, GRIDSS was applied to short read sequencing data from the NA12878 cell line (50X coverage PCR-free 2x100bp Illumina Platinum Genomes; ENA accession: ERA172924), along with several other structural variant callers. Callers were evaluated against both curated validated call sets [19, 30], and against PacBio and Moleculo long reads [31]. As previously [30], only deletions longer than 50bp were considered. So as to not unfairly penalise imprecise callers such as BreakDancer, calls were considered true positives if the breakpoint position error was less than the library fragment size, and the event length differed by at most 50% from the validated call set. For the PacBio and Moleculo long reads, variant calls required at least 3 split reads (with each split alignment mapping at least 25% of the long read), or 7 reads containing a corresponding indel, to support the event. ROC curves for other callers were obtained by varying the required number of supporting reads as reported by the caller.

For both validated call sets, GRIDSS exhibits generally-considerably better performance characteristics than other callers (Figure 3). When compared to the Mills *et al* call set, GRIDSS was able to identify the first 1,000 true positives with a false discovery rate of 7% (3% using PacBio/Moleculo validation data), compared to the next closest method, LUMPY, at 11% (7%). When an aligner that reports multiple alignment positions for each read is used, GRIDSS sensitivity is improved by 25%, but at cost of slightly reduced specificity (see Supplementary Materials for details). To determine the relative contributions of split read, read pair, and assembly support, GRIDSS was run on read pair (DP) and split read (SR) subsets with and without assembly. As expected, using more of the available evidence results in better variant calls (Figure 4). Assembly improves variant calling for DP and DP+SR evidence, but does not improve SR alone as the assembly contig lengths are limited to the length of the longest soft clip.

**Figure 3.**
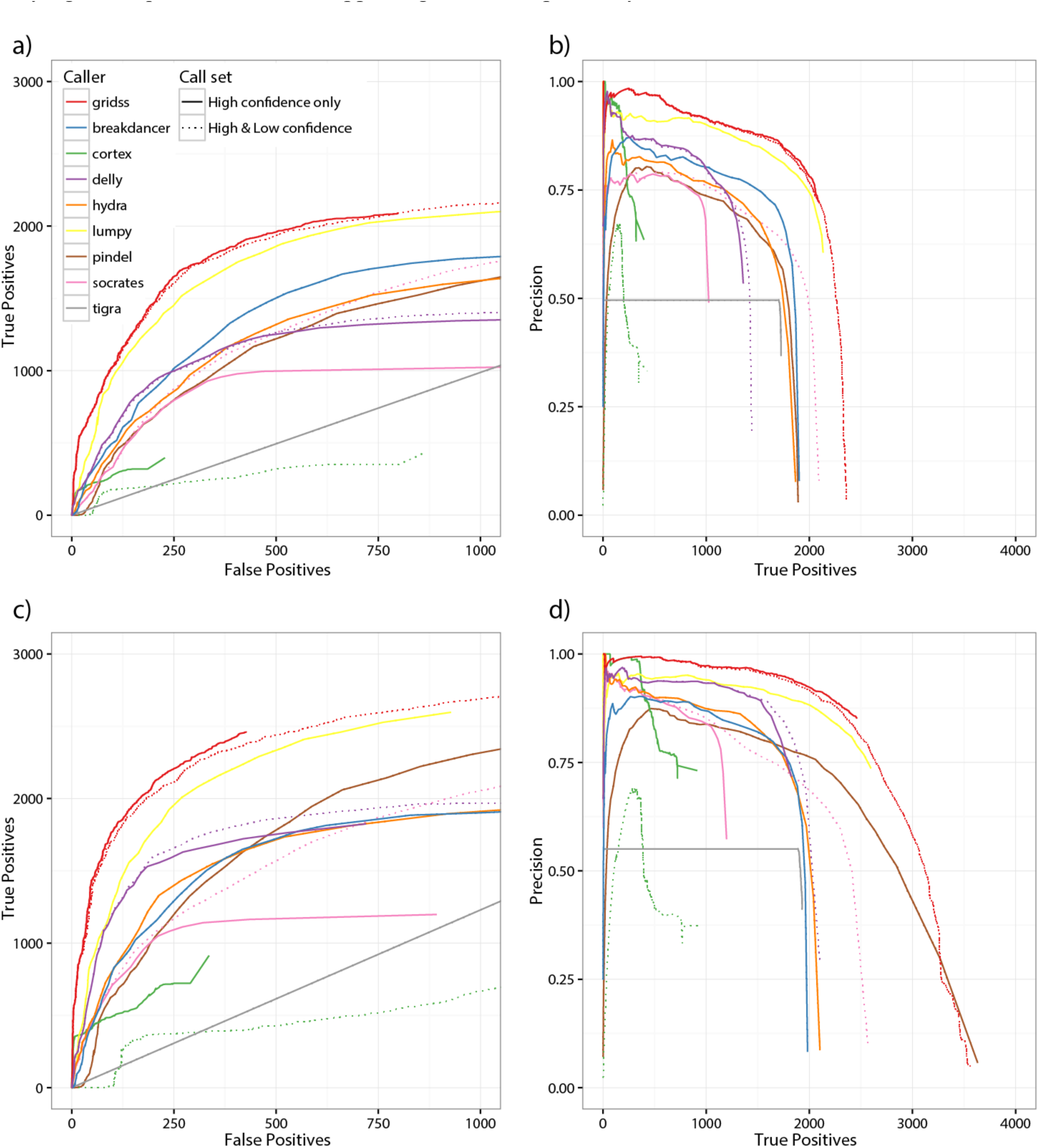
Performance of different SV callers on deletion detection in NA12878 at 50x coverage. Multiple variant calls were compared to both the Mills *et al* validation call set (a and b) and PacBio/Moleculo long read orthogonal validation data (c and d). Plots show the number of true positives versus false positives (a and c) and the precision versus true positives (b and d). PacBio/Moleculo validation required 3 split, or 7 spanning long reads supporting the call.

**Figure 4.**
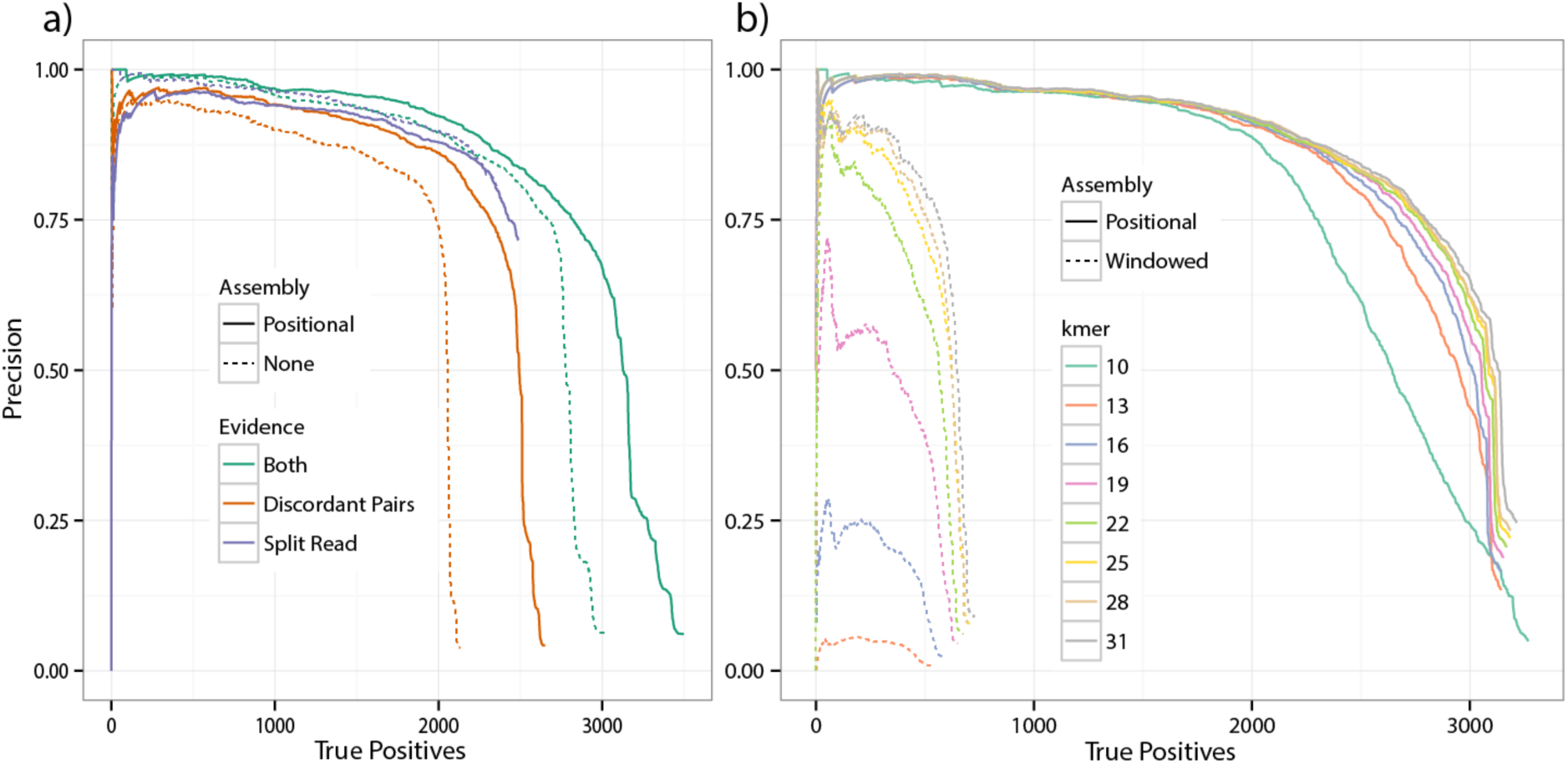
Performance of GRIDSS variant calling and assembly on NA12878 deletions events using PacBio/Moleculo orthogonal validation data. Precision versus the number of true positives for different types of support (a) and for different kmer sizes (b). Assembly of both split reads and read pairs improves both sensitivity and specificity to levels not achievable by either evidence source. Scoring only assembly-supported variants and varying the type of assembly and kmer size demonstrates that robust small kmer break-end assembly can be achieved with positional de Bruijn graph assembly, but not windowed de Bruijn assembly.

To understand the contribution of the positional de Bruijn graph assembly, we tested a windowed break-end assembler using a traditional de Bruijn graph (see Supplementary Material) in the GRIDSS framework. By restricting variant scoring to consider only assembly-based support, we compared windowed and positional assembly across a range of kmer sizes against the PacBio/Moleculo orthogonal validation data set. Positional de Bruijn graph assembly exhibited minimal drop in performance from kmers 31bp to 22bp, while windowed assembly performance was highly sensitive to kmer size, and sensitivity and specificity were well below those of the positional assembly for all kmers tested (Figure 4). The poor performance of windowed assembly was traced to the incorrect assembly of kmers occurring in multiple positions within the assembly window (Supplementary Figure 10). As expected, this phenomenon was especially pronounced in simple repeats and regions of low complexity (Supplementary Figure 7). As this mode of mis-assembly does not occur in positional de Bruijn graph assembly, GRIDSS is able to perform assembly with shorter kmers and therefore at lower coverage than either windowed or *de novo* assembly.

### Application to complex genomic rearrangements

To evaluate the performance of GRIDSS on complex genomic rearrangements, SVs were predicted in three published cancer-associated neochromosome datasets [32] (see Supplementary Materials). Each neochromosome contains hundreds of genomic rearrangements identified from the integration of copy number and discordantly aligned read pairs, followed by extensive manual curation. A high concordance with the 1010 curated SVs was obtained with GRIDSS detecting 98% of the curated calls, 92% with high confidence. GRIDSS obtained a higher concordance with the previously published curated results than other tested methods (BreakDancer, Cortex, DELLY, LUMPY, HYDRA, Pindel and SOCRATES; see Supplementary Materials).

GRIDSS also refined the original call set in three ways. Firstly, GRIDSS calls were made to single nucleotide resolution. Secondly, GRIDSS was able to identify 414 additional high confidence breakpoints, the majority (66%) of these were missing from the curated call set because they were supported by fewer read pairs than the fixed threshold applied in the original analysis, or were within 1000bp of another SV (potentially an issue due to the use of DP evidence alone). Finally, in 5% (64) of the SVs, GRIDSS was able to refine events classified as simple genomic fusions between two locations (A and B) that were in fact compound genomic fusions (from locations A to C to B), where the fragment C was short. In these events, GRIDSS was able to assemble a breakpoint contig at A, fully spanning the C fragment, with the remainder of the contig unambiguously aligning to B. A further 31 compound genomic fusions were identified in which the spanned fragment could not be unambiguously placed. While pure split read methods should also detect these compound rearrangements, the order in which DP and other evidence is applied and how it is applied will impact whether other methods can detect these features. This further refines the picture of complex rearrangements in neochromosomes and provides single nucleotide resolution of DNA breaks.

### Application to cancer samples

To demonstrate the clinical utility of the low false discovery rate of the GRIDSS high-confidence call set, GRIDSS was used to identify patient-specific DNA biomarkers from tumour biopsies for application to cell free DNA. Somatic rearrangements were predicted from 40x coverage sequencing data from primary lung cancers from two patients without matched germline data (see Supplementary Materials for details). Primers were designed for 8 candidate SVs from the GRIDSS high confidence call set (four from each patient) and all 8 SVs were validated by real-time PCR (only 6 were somatic, the remaining 2 were found to be real germline SVs).

Next, GRIDSS was compared to results from two previously published cancer data sets. First, SVs were identified from sequencing data from a melanoma metastasis and matched germline samples (60X coverage tumor; 30X coverage normal) [14]. GRIDSS detected 1,051,017 high and low confidence SVs with any level of support. Of these, 492 were high confidence calls and 851,981 had very low support (at most 3 supporting reads/read pairs). Of the 8 somatic events previously predicted by Socrates and validated by PCR [14], all 8 (100%) were identified by GRIDSS with high confidence.

Finally, GRIDSS was run on DNA-Seq data from the HCC1395 breast cancer cell line and results compared to the published genomic breakpoints, which were predicted to be associated with “validated” fusion genes [33]. GRIDSS showed strong concordance with the published results (Supplementary Table 2). Using a 21x coverage subset of the HCC1395 WGS data (only 21x of the 63x is publically available), GRIDSS identified 23 of the 26 published genomic breakpoints (22 with high confidence, 1 with low confidence), and failed to identify 1 breakpoint (manual inspection showed no supporting reads were present in the available data). The remaining 2 calls not identified by GRIDSS appear to be false positives caused by read-through transcription (see Supplementary Materials for details). Additionally, GRIDSS identified genomic breakpoints associated with a further 3 of the validated fusion genes.

### Application to *Plasmodium falciparum* genome data

To test the behaviour of GRIDSS on a challenging AT-rich non-mammalian genome, it was applied to a laboratory strain of *Plasmodium falciparum* that was genetically modified for use as a live, attenuated malaria vaccine (C5) and to the parental 3D7 population (ENA accession: PRJE12838). In the vaccine candidate, a plasmodium gene KAHRP was knocked out by insertion of a construct containing a processed copy of the human DHFR transcript. In addition to identifying the insertion of the construct into the KAHRP locus, GRIDSS detected 8 of the 9 exon splicing events associated with the processed DHFR transcript with high confidence, while the ninth splicing event was found in the low confidence call set. This demonstrates the value of GRIDSS’ sensitivity and prioritization of calls. Finally, a tandem duplication was also identified by GRIDSS in one of the VAR gene regions and supported by a copy number change (Figure 5). VAR genes are a large and complex gene family and the VAR gene regions are prone to recombination. The duplication was clonal in the C5 candidate and present at low frequency in the parental population (supported by 1 read), which was utilitized in the colored, positional de Bruijn graph to identify the rearrangement.

**Figure 5.**
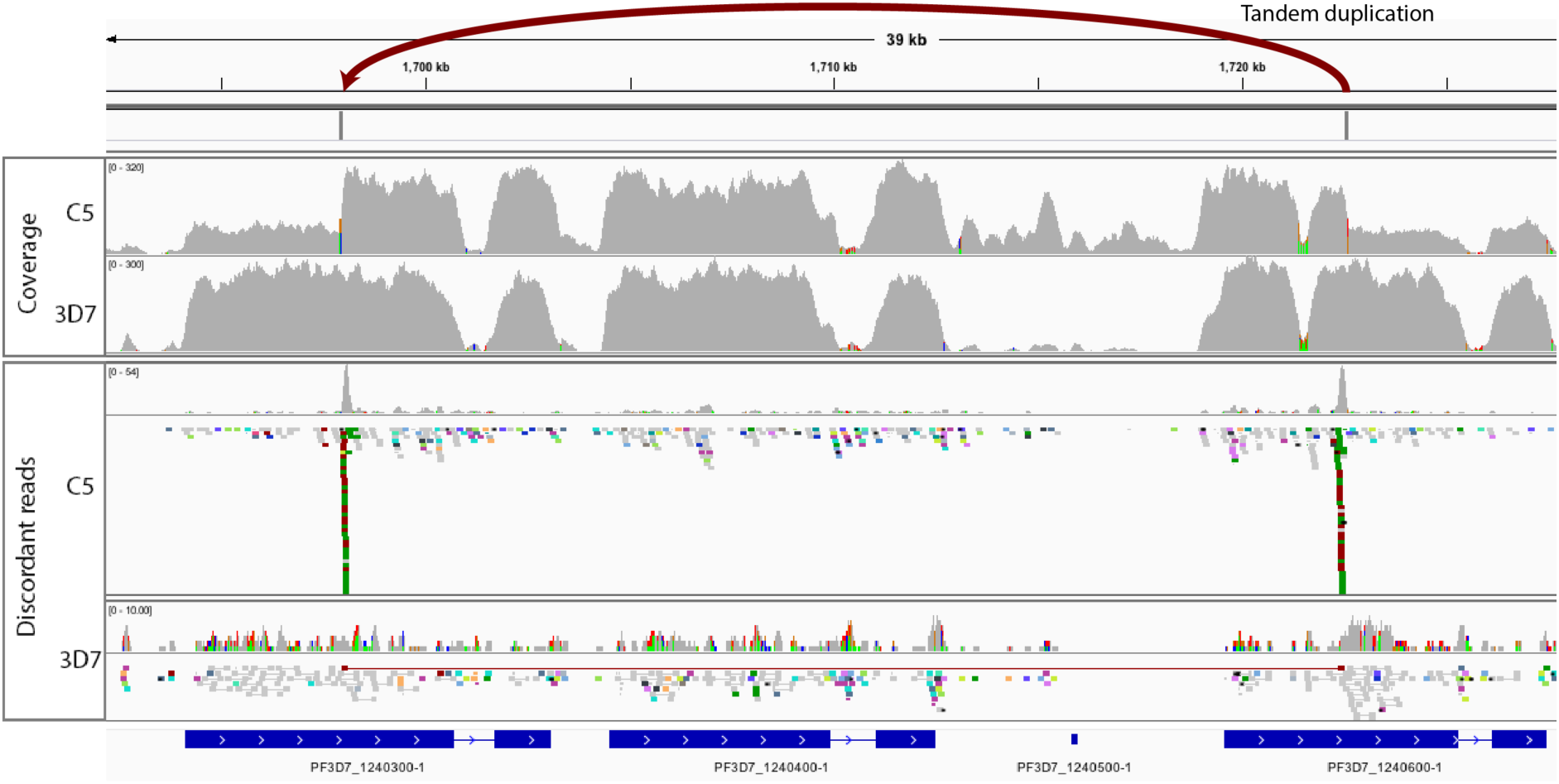
A tandem duplication identified in a VAR-gene region of the AT-rich *Plasmodium falciparum.* Coverage is shown for two samples of *P. falciparum*—a genetically modified line (C5), which was derived from the parental laboratory strain (3D7). The AT-rich genome shows high coverage in genes, which drops to very low levels in the AT-rich non-exonic regions. A change in copy number is apparent in the C5 coverage. GRIDSS detected the underlying tandem duplication in the C5 vaccine candidate (indicated). The supporting discordant read pair (DP) evidence is shown for both strains. Weak evidence (1 read pair) for this rearrangement was also detected in the parental population indicating that the SV was sub-clonal in this population. This evidence contributed to the coloured, positional de Bruijn graph assembly.

### Runtime performance

Runtime performance of variant callers was compared for 50x coverage whole genome human sequencing, and variant-dense simulated data sets on a virtual machine with 20 cores on a dual-socket Xeon E5-2690v3 server. GRIDSS runtime is comparable to that of LUMPY, lags behind that of BreakDancer, SOCRATES and HYDRA, and is considerably faster than DELLY, Pindel, Cortex and TIGRA (Table 1). Of the 236 minutes wall clock time taken by GRIDSS, 159 minutes was spent on file I/O and parsing of the input over three separate passes (library metrics calculation, evidence extract, reference coverage annotation), with only 38, 11, and 11 minutes spent on assembly, soft clip alignment by external aligner, and variant calling respectively (Supplementary Figure 15).

**Table 1.** SV caller runtime performance. Times shown are execution time in minutes. Benchmarking was performed on a dual socket Xeon E5-2690v3 server.

|  | SOCRATES | BreakDancer | HYDRA | LUMPY | GRIDSS | DELLY | Pindel | Cortex | TIGRA |
| --- | --- | --- | --- | --- | --- | --- | --- | --- | --- |
| NA12878 50x | 52 | 82 | 85 | 211 | 236 | 489 | 2184 | 8174 | 13196 |
| Simulation | 88 | 74 | 147 | 258 | 440 | 702 | 2475 | 17460 | 3660 |

## DISCUSSION

We have developed the GRIDSS software package, which performs genome-wide break-end assembly, and combines assembly, split read, and discordant read pair evidence using a probabilistic model to identify and score structural variants. It automatically detects non-template sequence insertions, and both exact micro-homologies and large imperfect homologies. Through comparison with existing split read, read pair, assembly-as-validation, and *de novo* assembly approaches on both simulated and real data, we have shown that our approach significantly improves both sensitivity and specificity of variant calling. We also demonstrated that GRIDSS is effective on real data from human tumours as well as non-mammalian organisms.

GRIDSS is designed to make the most of available evidence. The break-end assembly algorithm is able to make use of SR, DP, OEA, SC and indel-containing reads, and utilizes information generated by the aligner via a positional de Bruijn graph. By combining SR, DP and assemblies into a unified probabilistic model, GRIDSS is able to call variants even in the absence of any two of these signals. Such an approach is a distinct advantage over callers such as DELLY that require a threshold signal strength in one signal before considering any others.

GRIDSS is designed to be highly sensitive and not miss any putative genomic rearrangements. GRIDSS uses variant score and the presence of assembly support from both sides of the rearrangement to classify variants as high or low confidence. The high confidence set provides an immediately usable call set with high specificity, which is particularly important in clinical applications. The retention of low confidence calls enables analysis requiring high sensitivity.

Our novel genome-wide break-end assembly approach is made possible through the utilisation of a positional de Bruijn graph to incorporate read alignment constraints into the graph structure itself. This allows us to perform genome-wide assembly without targeting or windowing, and use a kmer size half that required for *de novo* de Bruijn graph assembly. This small kmer in turn results in improved assembly at lower levels of coverage. Although our approach results in a graph two orders of magnitude larger than the equivalent *de novo* assembly de Bruijn graph, we are able to perform genome-wide break-end assembly faster than a number of existing targeted breakpoint assembly implementations through graph compression, data streaming, multi-threading, reference-supporting read exclusion, and extensive use of dynamic programming and memoization. By demonstrating that GRIDSS performance is comparable to existing callers when only discordant read pairs or split reads are considered, we show that the incorporation of positional de Bruijn graph-based whole genome break-end assembly into the variant calling process is key to the superior performance of GRIDSS.

GRIDSS has been released as free and open source software under a GNU General Public License (GPL version 3) and is available at https://github.com/PapenfussLab/gridss.

## METHODS

GRIDSS has been designed for 36-300bp Illumina paired-end or single-end DNA-Seq sequencing data, and accepts any number of coordinate sorted SAM, BAM or CRAM input files, with no requirement for matching read lengths between input files. Mate-pair libraries are currently not supported. User-supplied categories for each input file allow for somatic and multi-sample calling with support and variant scoring broken down per category. Variant calls are output to VCF using the break-end notation of VCFv4.2.

### Identification of supporting reads

An initial parse of the read alignments is performed to collect library metrics. Metrics are calculated independently for each input file, thus allowing libraries from multiple related samples (or unrelated samples from a population) to be processed together. The following metrics are gathered in the initial pass for each input file: read alignment CIGAR element length distribution, fragment size distribution, and counts of total reads, mapped reads, unmapped reads, total read pairs, read pairs with both reads unmapped, read pairs with one read unmapped, read pairs with both read mapped, maximum read length and maximum mapped read length. The fragment size distribution is calculated using Picard tools CollectInsertSizeMetrics.

Reads partially aligning to the reference (reads containing indels or soft clipped bases) and read pairs in which the inferred fragment size falls outside the expected fragment size distribution (default shorter or longer than 99.5% of read pairs) of the library, with incorrect orientation, or with only one read mapped are extracted to intermediate files. GRIDSS does not utilise read pairs in which both reads are unmapped.

### Split read identification

Split reads are identified by aligning the soft clipped bases of partially aligned reads back to the reference using bwa (default) or another aligner that is compliant with the SAM file format specifications (such as bowtie2). Except for the case where multi-mapping alignment is used, reads for which the soft clipped bases are uniquely aligned to the reference (default MAPQ ≥10) are considered to provide split read support. Reads containing insertions or deletions in the read alignment are treated as split reads aligning to either side of the indel.

### Read filtering

Due to the prevalence of artefacts in sequencing data, GRIDSS includes a number of default filters designed to improve specificity by removing reads likely to interfere with variant calling. By default, the following read filters are applied:

- MAPQ at least 10. The read mapping quality score (MAPQ) is defined by the SAM specifications as −10 log_10_(Pr(incorrect mapping)). A low mapping quality score indicates that the read is unlikely to be mapped correctly. All reads with a MAPQ score of less than 10 are treated as unmapped. For split reads, both the read MAPQ and the soft clip re-alignment MAPQ must exceed the threshold.
- Maximum coverage of 10,000x. Regions of extremely high coverage may be due to a collapsed reference sequence in highly repetitive regions. Coverage of canonical sequences such as those present in the centromeres can be greater than 3 orders of magnitude higher than the nominal coverage and as such, regions with coverage greater than 10,000x are excluded from variant calling and assembly.
- Soft clipped reads must have at least 95% of aligned bases matching the reference sequence as these partially aligned, poorly matching reads are frequently alignment artefacts.
- Soft clipped reads contain at least 4 soft clipped bases. Sequencing errors in the bases near either end of a read will result in Smith-Waterman local alignment of the read excluding the end bases from the alignment.
- Estimated Shannon entropy of aligned nucleotides (based on frequencies of nucleotides in the read) of at least 0.5 bits. Reads with low complexity in the mapped bases (such as poly-A runs) were found to have a high false positive rate. If the estimated entropy of nucleotide frequency distribution within the read is less than 0.5 bits, the read is considered unmapped. For example, a read with sequence AAAAAAAAAGTTT with the final three bases soft clipped would have estimated entropy of 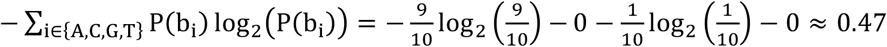 bits and would be considered unmapped.
- Mean Phred-scaled base quality score of soft clipped bases must be at least 5. Soft clips containing runs of low base quality scores are filtered as such sequences are typically due to sequencing errors.
- Adapter filtering. Reads containing the first 12 bases of the Illumina Universal Adapter, Illumina Small RNA Adapter, or Nextera Transposase sequences are filtered if the sequence is homologous to the reference sequence at the aligned location.
- Overlapping read pairs from short fragments are excluded. A read pair aligned in the expected orientation with overlapping read alignments is indicative of pair sequenced from a fragment shorter the combined read lengths. Such read pairs do not provide support for structural variants and are filtered. If soft clipped, each read is still individually considered for split read analysis and assembly. In the case that the fragment length is shorter than the read length, the reads may contain soft clips ‘dovetailing’ in adapter sequence past the other read. If there exists a micro-homology between the adapter and reference adjacent to the alignment position, the reads will be over-aligned and such read pairs will appear to be in an unexpected orientation. Such edge cases are filtered by allowing a 4bp dovetail when determining whether a read originates from a short fragment.

### Positional de Bruijn graph assembly

A positional de Bruijn graph is a graph where each node contains a sequence of k bases and the position at which those k bases are expected to occur. The positional de Bruijn graph *G* = (*V, E*) consists of the vertex set *V* and the edge set *E* ⊆ *V×V*. Let *n* = (*S_n_,p_n_*) ∈ *V* where *S_n_ =* (*b*_1_*, b*_2_,…, *b_k_*) is a kmer of *k* bases, and *p_n_* ∈ ℤ is the expected genomic position of the kmer. By definition, nodes are connected if they have adjacent kmers and positions:

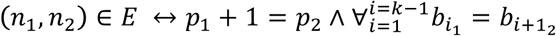

Unlike a traditional de Bruijn graph, *G* is a directed acyclic graph since ∀*_i_,_j_*(*n_i_, n_j_*) ∈ *E* → *p_i_* < *p_j_*; any cycle would contain an edge in which *p_i_* ≥ *p_j_*. That is, all paths through the graph are simple (not self-intersecting) paths since traversal of every edge must advance the position by a single base. This allows algorithms such as the longest path problem that would require exponential time on a de Bruijn graph to be completed in polynomial time when applied to a positional de Bruijn graph. In addition to the kmer and position, the following node attributes are used:

- *w*(*n*) is the node weight corresponding to the Phred-scaled probability of the supporting read kmers. The probability of the *i*th kmer of read *r* is given as the joint probability of every kmer base being correctly called by the sequencer, and the aligner mapping the read to the correct location. That is,

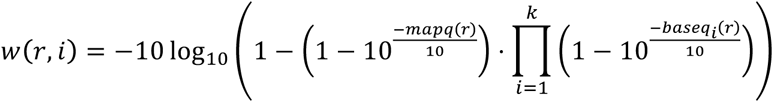

where *baseq_i_* (*r*) is the Phred-scale base quality score of the *i*th base, and *mapq*(*r*) is the mapping quality score of read *r*. As *w*(*r, i*) is Phred-scaled, *w*(*n*) = Σ *w*(*r, i*).
- *anchored*(*n*) is a boolean flag indicating whether the node bases are fully aligned to the reference genome in any supporting read. Note that this definition does not require the read bases to actually match the reference and both matched and mismatched base alignments are considered to be anchored if aligned.
- *Support*(*n*) is the set of reads providing support for the given kmer at the given position.

### Assembly graph construction

All extracted reads are added to one of two positional de Bruijn graphs based the expected break-end orientation. Reads supporting a rearrangement after the anchored position are added to the forward graph, whereas reads supporting a rearrangement before the anchored positions are added to the backward graph. Each read kmer is added to the graph at all expected mapping positions. For a soft clipped read, each read kmer is added to *G* at the position the kmer would start at if the entire read was mapped to the reference. Split reads are treated as two independent soft clipped reads. For read pairs, read kmers are added at all positions in which the read pair would be considered concordantly mapped based on the mapping location of the partner. For discordant read pairs, each read is added based on the anchoring location of the mate irrespective of the actual mapping location of the read, whereas one-end anchors are only added at one location as the unmapped read can provide no positional constraints on its partner.

Using this method of graph construction, correctly assembled unanchored paths are limited in length to less than the read length if supported by only soft clipped or split read evidence, and by the maximum concordant fragment size if supported by read pair evidence.

### Assembly graph error correction

Base calling error correction is performed by collapsing similar paths. Paths are scored according to the sum of the node weights. A path A is collapsed into an alternate path B if both the total path weight of A is less than B, A and B differ by less than a fixed number of bases (default 2), and either both paths are same length and share the same start and end node (bubble popping), or A shares a start or end node with B and contains a single terminal leaf node (leaf collapse). By default, error correction is only performed on paths of length less than twice the read length that are either simple bubbles or terminal leaves.

For each node, all leaf and branch paths under the maximum length originating from the node are identified by traversal of branchless descendants. For each path identified, breadth first graph traversal is performed to identify candidate paths to merge. Memoization is used to track the optimal paths to each node thus reducing worst-case traversal complexity from exponential time to quadratic.

### Break-end contig assembly

A break-end contig path consists of a sequence of adjacent unanchored nodes, optionally flanked by a sequence of anchored nodes. Paths flanked by anchoring nodes are called before unflanked paths. Maximally weighted paths are calculated in the same manner as error correction traversal with breadth-first traversal with memoization of the highest-weighted partial path at each node.

Once the maximally weighted path has been determined, the path is extended into flanking anchored nodes until the anchored path length exceeds both the maximum read length and the unanchored path length. Once the maximally weighted path contig is called, all reads supporting any unanchored kmer on the contig path are removed from the graph. The removal of supporting reads from all graph nodes ensures that each read contributes to a single assembly only. Contigs are iteratively called until no non-reference kmers remain in the graph.

Assembly contigs supported by less than the minimum required support (by default 3 reads) and unanchored contigs shorter than the read length are filtered.

### Contig error correction

While positional information at nodes significantly reduces the rate of mis-assembly compared to windowed assembly, branch traversal introduces new modes of mis-assembly. When a read pair is self-intersecting or contains repeated kmers, the resultant contig will loop for as long as the fragment size window will allow. This mis-assembly can occur even with single read. For example, with k=4, the single read TAAAAC expected to start at one of the positions in the interval [10, 15], will result in the highest weighted path of TAAAAAAAAC starting at position 10. To prevent such mis-assemblies, kmer chaining of the supporting reads is used to truncate the called path at the first kmer transition not supported by any constituent read. Truncation is performed starting from both the start and the end of the contig with the highest weighted truncated path called, preferentially calling anchored paths. Each supporting read is aligned to the contig position with the greatest number of matching kmers (breaking ties towards the truncation start kmer) and all kmer transitions supporting by the read are marked. Once all supporting reads have processed, the contig is truncated at the first kmer transition not supported by any reads.

### Contig re-alignment

Once an assembly contig has been called, a multi-stage re-alignment process is used to identify the breakpoint supported by the contig. Assemblies containing at least one anchored base undergo Smith-Waterman (local) re-alignment around the expected contig position. Assemblies that fully align to the reference are treated as a fully-aligned indel-spanning assembly if an indel is present in the alignment, or filtered out as a false positive if the full alignment contains no indels. For unanchored assemblies with no soft clip or split read support, a breakpoint position interval is calculated based on the breakpoint interval consistent with the greatest number of supporting read pairs.

The contig bases not anchored to the reference are aligned using an external aligner (by default bowtie2) in local alignment mode using the same alignment thresholds used for identifying split reads. For assembly alignment, if the external aligner identifies the best alignment to be a soft clipped alignment, these soft clipped bases are again aligned, with such recursive alignment limited to a depth of 4. This compound re-alignment results in assembly support not only for the breakpoint site identified by the initial re-alignment, but also for any additional breakpoints spanned by the assembly. This approach allows accurate classification of complex rearrangements such as those present in neochromosomes formed through chromothripsis and the breakage-fusion-bridge process.

### Detection of micro-homology and non-template sequence insertions

Depending on the pathway involved in DNA repair resulting in a structural variant, there may exist either sequence homology at the breakpoints, or non-template sequence inserted during the repair. Micro-homology at the breakpoints introduces uncertainty into the breakpoint position call, which is resolved if non-template sequence is also present. As a consequence of aligner behaviour, unhandled sequence homology can result in two separate variant calls (one at each edge of the homology) for a single event as aligners are able to map read to the homologous sequence at both side of the breakpoint. Unhandled non-template sequence insertions results in incorrect breakpoint sequence and event size calculation and, for coding variants, result in incorrect gene fusion transcript prediction. GRIDSS factors in both breakpoint sequence micro-homology and non-template sequence insertions when performing variant calling. For micro-homologies, the breakpoint location for split reads and assemblies is expanded from a single base to an interval of the length of sequence homology between the read/assembly and the reference sequence at either side of the predicted genomic rearrangement. The nominal position of the called breakpoint is considered to be the centre of the homology, and is reported using the standard HOMSEQ, HOMLEN, and CIPOS fields in the VCF output. Non-template sequence insertions are included as an additional component of the variant call in a similar fashion to SOCRATES [14].

In addition to the standard VCF homology fields, GRIDSS reports the inexact homology in the nonstandard IHOMPOS field. The inexact homology length is calculated for each break-end by performing local Smith-Waterman alignment of the breakpoint sequence to the reference sequence up to 300bp either side of the break-end.

### Graph compression

To reduce the graph size, a compressed representation of the positional de Bruijn graph is used. Nodes with a single successor connected to nodes with a single ancestor correspond to a branchless path and are compressed into a single path node. Similarly, nodes with adjacent positions and matching kmers, weights and reference status are compressed using a position interval. The resultant path nodes are of the form (*start, end*, (*kmer*_1_*,weight*_1_,… *,kmer_n_,weight_n_*)*, anchored*), representing the set of nodes 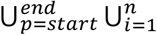 (*p, kmer_i_, weight_i_ anchored*).

### Streaming assembly

Since all nodes are associated with a genomic position and only have edges to adjacent genomic positions, all graph operations only affect the local subgraph near the genomic position. Exploiting this, positional de Bruijn graph assembly is performed in a single streaming pass over the input reads. Coordinate sorted records are streamed through the following 5 internal processes, each processing records within a genomic position window size determined by the maximum read length and fragment size:

1. Extraction: each read is converted into constituent positional de Bruijn graph nodes. To reduce the graph size, positional intervals are stored implicitly with each extracted node defined over a positional interval.
2. Positional aggregation: overlapping nodes from multiple reads are split into disjoint aggregated nodes and graph edges calculated and cached.
3. Path compression: non-branching aggregate node paths are compressed into path nodes.
4. Error correction: bubble popping and leaf collapse error correction is performed to remove base calling artefacts.
5. Assembly: maximal path contig calling is performed such that whenever a maximal path is encountered, the streamed assembly graph loads and traverses all alternate paths which any reads supporting the maximal path could also support. Since each read can contribute to graph node positions over an interval no wider than the concordant fragment size plus the read length, the globally maximal path containing any given read must overlap this interval and is thus a local graph operation. A contig is called whenever the subgraph has loaded all potential alternate paths for the highest weighted maximal path encountered. Since all reads contributing to the called path must have been fully loaded for all such potential alternate paths to have been traversed, all reads contributing to the contig are fully removed from the subgraph immediately. This approach ensures that the any contig called is the globally maximal contig containing the given reads.

This assembly algorithm is implemented by memoization of all maximal paths of the streaming subgraph. The starting position of paths for which a potential successor node has not yet been loaded are tracked in a frontier and when the end position of the maximal path plus the concordant fragment size is earlier than both the earliest start position of the frontier path and the start position of the next node to be loaded, the maximal contig is called.

As a full recalculation of all maximal paths in the subgraph after supporting reads have been deleted is unnecessary, only paths affected by node removal are recalculated. For sufficiently high-coverage data, there will be enough concordant fragments with unexpectedly long or short fragment size that a background signal supporting small indels everywhere across the entire genome will be present. When these reads are assembled, this signal results in long unanchored paths up to multiple megabases in size. To supress this background noise, contigs and path longer than the maximum expected size are filtered and thresholds are placed on both the size and the genomic width of the loaded subgraph (See Supplementary Materials).

### Coverage blacklist

Alignment of whole genome sequencing data to the reference results in extreme coverage in some genomic regions, for example in satellite repeats. These regions are likely to contain false positives and are computationally expensive to assemble due to the high coverage. To mitigate the effect of such regions on variant calling and runtime performance, GRIDSS performs two independent operations. Firstly, GRIDSS filters all reads within 1 fragment length of any region of extreme coverage (default 10,000x). Secondly, GRIDSS down-samples before assembly in poorly mapped regions, where the density of variant-supporting reads is unusually high. When the number of SV-supporting reads within a sliding window (default 1,000bp) exceeds the threshold (default 5,000), only a subset of the reads are used for assembly. The probability of a SV-supporting read being retained is given by

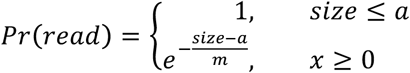

where *m* is the target maximum number of reads in the window, and *a* is the unconditional acceptable threshold in which no downsampling occurs (default 50% of maximum). This hybrid exponential back-off model ensures that no down-sampling is performed except for coverage outliers, and within coverage outliers, real events are unlikely to be filtered.

### Variant calling

#### Probabilistic variant scoring model

To estimate the quality of predicted structural variants, we score variants according to the Phred-scaled probability of originating from the mapped locations without any underlying structural variations. The Phred score *Q* of a probability *P* is given by *Q* = −10 log_10_ (*P*). Given a read pair or split read *r* mapping to genomic locations *a* and *b,* and the event *M* that the mapping is correct, the score assigned to *r* is given by Pr(*r*) = Pr (*r*∩*M*) *+* Pr(*r*∩*M̅*) = Pr(*M*) ⋅ Pr(*r|M*) + Pr(*M̅*) ⋅ Pr(*r|M̅*). The probability of correct mapping Pr(*M*) is determined directly from the Phred-scaled mapping quality scores *mapq_a_*(*r*)*, mapq_b_*(*r*) defining the probability of incorrect read alignment: Pr(*M*) = (1 − 10^*mapq_a_*(*r*)/10^)(1 − 10^*mapq_b_*(*r*)/10^) = 1 − Pr(*M̅*) since Pr(*M*) requires both mapping locations to be correct. In the case of incorrect mapping, *r* is uninformative and Pr(*r*|*M̅*) = 1. In the case of correct mapping, Pr(*r|M*) is determined empirically from the relevant library distribution.

For split reads, correct mapping with no SV implies the alignment is artefactual. We model the artefactual alignment rate from the empirical soft clip length distribution of the library thus Pr(*r*|*M*) = *P_sc_*(*l_sc_*(*r*)) where *l_sc_*(*r*) is the length of the soft clip of *r* before split read identification, and *P_sc_* is the library soft clip length distribution.

For read pairs, correct mapping can be caused either by a chimeric fragment *CF,* or an unexpectedly large/small originating fragment: Pr(*r*|*M*) = Pr(*r* ∩ *CF|M*) + Pr(*r* ∩ *C̅F̅|M*) = Pr(*CF*|*M*) ⋅ Pr(*r|M* ∩ *CF*) *+* Pr(*C̅F̅*|*M*) ⋅ Pr(*r|M* ∩ *C̅F̅*). *Pr(r\M* ∩ *C̅F̅*) is given as *P_rp_*(*ifs*(*r*)) where *ifs*(*r*) is the fragment size of *r* inferred from the read mapping locations, and *P_rp_* is determined from the library fragment size distribution inferred from all mapped read pairs. Chimeric fragment alignments are considered uninformative and *Pr*(*CF|M*) is taken to be *p_d_*, the rate of read pair mapping in which the inferred fragment size exceeds more than 10 mean absolute deviations from the median library fragment size. Thus for read pairs Pr(*r|M*) *= p_d_ +* (1 − *p_d_*) *∙ P_rp_*(*ifs(r*)).

Split reads originating from indels use the corresponding distribution of insertion or deletion alignment operations instead of the soft clip distribution. Assemblies are modelled as a set of constituent reads with the anchored mapping quality defined as the greatest mapping quality of the constituent reads, and unanchored mapping quality determined by the assembly alignment mapping quality. Constituent soft clipped reads and reads with unmapped mates are treated as split reads and discordant read pairs for the purpose of determining assembly quality with the caveat that reads with unmapped mates use *p_u_* in place of *p*_*d*_. This assembly scoring model improves variant calling by rescuing poorly mapped reads, increasing the score of variants supported by assembly, and promoting SC and UM reads to SR and DP reads within the context of the assembly thus allowing these reads to be used as input to the variant calling. For computational efficiency, scoring calculations are approximated using the maximum Phred score of the constituent terms.

#### Variant calling using maximal cliques

Variants are scored according to the level of support provided by SR, DP and assembly evidence combined. Supporting evidence can be summarised as the tuple (*s_l_,e_l_,s_h_,e_h_,d_l_,d_h_,w*) where the intervals [*s_l_*, *e_l_*] and [*s_h_*, *e_h_*] are the genome intervals between which a breakpoint is supported; *d_l_* and *d_h_*, the direction of the supported breakpoint; and *w* the weight of the evidence as defined by the evidence scoring model. Since each piece of supporting evidence is considered to be independent, and evidence scores are expressed as Phred scores, the score for any given variant is equal the sum of the scores of evidence supporting the variant breakpoint.

Calculating the total support weight for all putative breakpoints is equivalent to finding all maximum evidence cliques, that is, all sets of evidence providing consistent support for a breakpoint that no more evidence can be added and the set remain consistent. Since both direction *d_l_* and *d_h_* must match if evidence is to mutually support a breakpoint, the evidence set can be divided into the four ++, +−, − +, −− directional subsets which reduces the problem to weighted maximum clique enumeration of a rectangle graph [Lee 1983]. Maximal clique enumeration is performed in a single in-order pass over evidence in polynomial time.

Unfortunately, this approach can result in reads providing support to multiple independent breakpoints. Since each read will have originated from at most one of the competing explanatory variants, a second pass is made and a greedy assignment is performed in which each piece of evidence assigned to only the highest scoring variant it supports.

#### Breakpoint position error margin

Split reads originating from the same event should only support a single breakpoint position. In practice, however, sequencing errors, imperfect homologies in the neighbourhood of the breakpoint, and alignment artefacts can result in variation of the breakpoint position supported by each read. As a result, split reads originating from the same event can provide support for breakpoints at different positions. To ensure such evidence results in a single call, an additional margin is added when determining clique assignment. By default, this margin is 10bp for variants 20bp or larger, linearly decreasing to no margin for 1bp thus ensuring that small non-overlapping indels are not merged.

#### Software development methodology

GRIDSS was developed as professional quality software using a test driven development methodology. To develop new functionality, test cases were first written describing the expected behaviour under normal and error conditions. Once such failing test cases have been written, code implementing the feature is written thus ensuring that the feature is functioning as expected. Bug fixing is performed by first creating a failing test case reproducing the error, then updating the implementation to correct the error. As a result, an extensive test suite composing of over 1,200 test cases has been developed. Git is used as a version control system. All tests are rerun prior to each release, ensuring regression faults in existing functionality are not introduced when new features are added.

Bug fixes and enhancement in libraries used by GRIDSS have been contributed back to these upstream libraries. Maven2 is used for build packaging and a single precompiled binary including all dependencies (except the external aligner) is produced for each release. All parameters used by GRIDSS (including the choice of external aligner used) have been externalised into a configuration file able to be modified by advanced users. Semantic versioning is used for release versioning. GRIDSS is implemented in Java 1.8. GRIDSS has been designed as a modular software suite. Although an all-in-one entry point is included, each stage of the GRIDSS pipeline, including the break-end assembler, can be run as an independent program, or replaced with an equivalent implementation. Example scripts for single sample, somatic, and multi-sample pipelines are provided. Java utility programs and R scripts to convert GRIDSS VCF break-end format to more user-friendly formats for downstream filtering and analysis are also provided. All scripts required for independent replication of results presented in this paper are included in the GRIDSS source code.

## DATA ACCESS

GRIDSS has been released as free and open source software under a GNU General Public License (GPL version 3) and is available at https://github.com/PapenfussLab/gridss. All scripts required for independent replication of results presented in this paper are included in the GRIDSS source code.

## ACKNOWLEDGMENTS

A.T.P. and T.P.S. were supported by an Australian National Health and Medical Research Council (NHMRC) Program Grant (1054618). D.C. was supported by an Australian Postgraduate Award. The research benefitted by support from the Victorian State Government Operational Infrastructure Support and Australian Government NHMRC Independent Research Institute Infrastructure Support. We thank Alan Rubin for helpful discussion on algorithm design.

## DISCLOSURE DECLARATION

The authors declare that they have no competing interests.

